# The impact of a governmental cash transfer programme on tuberculosis cure rate in Brazil: A quasi-experimental approach

**DOI:** 10.1101/311589

**Authors:** Daniel J. Carter, Rhian Daniel, Ana W. Torrens, Mauro N. Sanchez, Ethel L. N. Maciel, Patricia Bartholomay, Draurio C. Barreira, Davide Rasella, Maurício L. Barreto, Laura C. Rodrigues, Delia Boccia

## Abstract

**Background:** Social vulnerability is strongly associated with tuberculosis (TB) indicators like cure rate. By addressing key social determinants, social protection policies such as Brazil’s Bolsa Família Programme (BFP), a governmental conditional cash transfer, may play a role in TB control. Evidence is consolidating around a positive effect of social protection on TB outcomes, however methodological limitations prevent strong conclusions. This paper uses a quasi-experimental approach to more rigorously evaluate the effect of BFP on TB cure rate.

**Methods & Findings:** The data source was Brazil’s TB notification system (SINAN), linked to the national registry of those in poverty (CadUnico) and the BFP payroll. Propensity scores (PSs) were estimated from a complete-case logistic regression using covariates from this linked dataset, informed by a directed acyclic graph. Control patients were matched to exposed patients on the PS and the average effect of treatment on the treated (ATT) was estimated as the difference in TB cure rate between matched groups (n = 2167). The ATT was estimated as 10·58 (95% CIs: 4·39, 16·77). This suggests that 10·58% of the TB patients receiving BFP who were cured would not have been cured had they not received BFP. The direction of this effect was robust to sensitivity analyses performed and the PS matching broadly improved balance, although missing data limited the sample size.

**Conclusions:** This work is the first quasi-experimental evaluation of social protection in wide-scale practice on TB outcomes. It demonstrates a positive effect of conditional cash transfers on TB cure rate consistent with existing work, suggesting changes to policy and future research on increasing access to social protection for TB patients who remain uncovered by the programme.

## Introduction

Despite biomedical efforts, the global burden of Tuberculosis (TB) remains considerable, with up to 1.5 million deaths from TB recorded in 2015 [1]. Tuberculosis treatment takes many months, and a proportion of patients are not cured, either because they abandon treatment, take treatment irregularly, are infected with drug resistant TB, or die before completion of treatment [1]. The correlation between TB indicators and global poverty has been demonstrated both at ecological and individual level, yet much of the morbidity and mortality in TB patients still occurs amongst the poorest segments of the population [2]. Social determinants impact vulnerability to TB at every stage of the disease pathway, from TB infection to clinical outcomes, including whether or not the patient was cured [3]. Ending the global burden of TB requires bold policies and supportive systems able to recognise and tackle these social determinants [4].

Recognising this social aspect of TB epidemiology, social protection is now a non-negotiable component of the global response to TB, which also reflects the perfect alignment between the post-2015 END TB strategy and the Sustainable Development Goals agenda [4,5]. Brazil in particular has been an early adopter of this new policy, as reflected by its long term efforts to integrate development and health agendas. This is partially due to the strong social protection tradition in Latin America, which in Brazil culminated with the creation of the Bolsa Família Programme in 2003, one of the largest conditional cash transfer programmes in the world [6].

In 2010, the Bolsa Família Programme (BFP) provided a variable monthly stipend to households meeting certain socioeconomic criteria: households earning less than 70 reais a month (~22 USD at time of writing) and households with children, adolescents, or pregnant women earning less than 140 reais a month. BFP’s targeting is not exact, and individuals reporting an income above 140 reais can be found in the BFP payroll.[6] In order to receive BFP, families must be registered in the Cadastro Unico (Single Registry; CadÚnico), a registry of all low income Brazilian families. In return for the transfers, recipients must comply with behavioural obligations (i.e. school attendance; immunization). BFP is not explicitly intended to target TB-affected households and only ¼ of TB patients in Brazil are enrolled in the programme; given the intimate association between poverty and TB, underenrolment is likely [7].

Albeit accumulating, the literature on the impact of conditional cash transfers on a variety of TB indicators is still limited, and there has been little methodologically rigorous evaluation of social protection interventions for TB prevention, care, and control, including treatment outcomes. There has been some non-experimental support for cash transfers as cash incentives for TB, but we consider this work to come from a separate evidence base as the underlying philosophy and mechanisms of action for cash incentives differ greatly from cash transfers as social protection.[8] Despite its scarcity, the evidence is converging upon a consistent positive impact of social protection on TB epidemiology and control, including some small scale trials and studies in Peru and Moldova, an ecological study in Brazil, and a recent study undertaken in Rio de Janeiro [9–11]. Another recent study that encompassed the whole of Brazil, Torrens et al. (2016), showed that TB patients enrolled in BFP were approximately 7% more likely to be successfully cured after treatment than a control group [7]. Prior studies may have used control groups resulting in a biased comparison.

For an unbiased estimate of the proportion of patients cured attributable to BFP, we must construct a control group as similar as possible to the group of BFP recipients. This group of BFP recipients on average have some TB cure rate. We wish to estimate the difference in that cure rate if, counter to fact, that group of patients had not received BFP, but had the same sociodemographic characteristics and were thus still enrolled in CadÚnico.

To this aim, we approach the same routine data source as in Torrens et al. (2016) using a quasi-experimental approach to construct a more appropriate control group and to then determine a more rigorous estimate of the effect of BFP on TB cure rate amongst those who receive it. Specifically, we aimed to: 1) use propensity score matching to create a control group balanced for propensity to receive BFP, 2) provide an estimate of the average treatment effect of BFP on TB cure rate amongst recipients and 3) to reflect on the utility of the resulting estimate for changing TB policy.

## Methods

### Conceptual Framework - DAG

A directed acyclic graph (DAG) was proposed for conceiving of the causal relationships between the outcome, the exposure, and all the variables hypothesised to be on the causal pathway (Figure 1). Each node in the DAG consists of a high-level construct measured by proxy variables taken from the set of covariates available. The nodes in this DAG were constructed based on a variety of theoretical literature, and the grouping of covariates under one node denotes that they are considered to be measures of that underlying construct for the purposes of this paper [3,12–14]. Appendix 1 outlines explicitly which covariates fall under each node.

**Fig 1.**
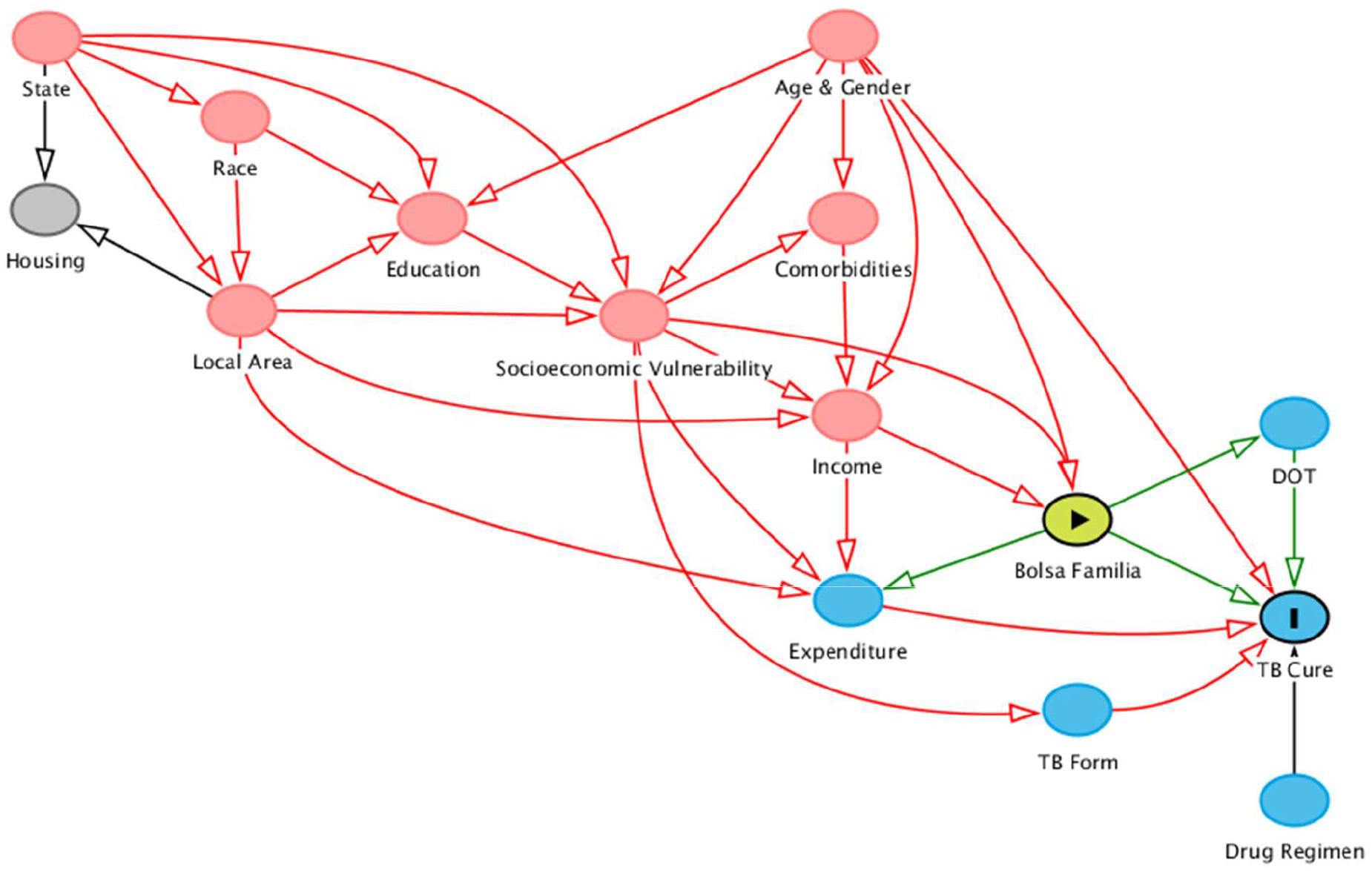
DAG for BFP and TB. A directed acyclic graph (DAG) built to conceptualise the potentially causal relationships between constructs relevant for measuring the impact of Bolsa Familia on TB Cure Rate. The DAG links nodes that represent constructs and are measured by covariates. These nodes include *Sex & Age* (Sex, Age), *Race* (Race, Indigenous, Quilombola), *Education* (Education Level, Literacy), *Local Area* (Urban, Running Water, Sewage, Electricity, Water Store, Trash), *Type of TB* (Thorax X-Ray, Initial Bacilloscopy, Pulmonary/Extrapulmonary, Throat Culture, Tuberculin Skin Test), *Directly Observed Treatment* (DOT), *Drugs* (Rifampicin, Isoniazid, Ethambutol, Streptomycin, Pyrazinamide, Ethionamide, Other Drugs), *Comorbidities* (AIDS, Alcoholism, Diabetes, HIV, Mental Disorder, Other Disorder), *Expenditure* (on Food, Energy, Gas, and Water), *Social Vulnerability* (Child Worker, Institutionalised, Work Acquired TB), *State*, and *Income*.

The DAG outlines potential mechanisms by which BFP (“the exposure”) is proposed to affect cure rate (“the outcome”). These include via access to directly observed treatment and via increased capacity for mitigation of catastrophic costs (expenditure). We provide an estimate for the direct effect of social protection outside of these pathways, which may include expanded access to healthcare through means other than DOT, increased psychosocial wellbeing, or greater integration into governmental systems in general. The DAG also outlines pathways between cure rate and income (and therefore access to BFP), through complex relationships between demographics, geography, and socioeconomic factors. The ‘cure’ outcome also includes those who completed treatment without bacteriological confirmation.

### Data Handling

The data for this study arose from a linkage between the 2010 TB dataset from SINAN (Brazil’s national Notifiable Disease Surveillance System) and the 2011 CadÚnico dataset The CadÚnico dataset was itself linked to the Bolsa Familia payroll held by the Caixa Federal (Federal Bank). The linkage added the demographic and social information from CadÚnico and the BFP payroll to every TB patient in the SINAN dataset.

Of the complete SINAN-CadÚnico-BFP dataset (n = 180046), only individuals who were new TB cases registered in CadUnico in 2010 with a non-missing treatment outcome variable were retained for this study (n = 16760). Exposed individuals (those receiving BFP) were further restricted to those whose receipt of BFP preceded case closure. Case closure is defined as the date on which an outcome (e.g. cure, unsuccessful completion of treatment, death) is recorded. The final dataset used for analysis included 13,029 individuals, 6,940 of whom received BFP.

The dataset contained a set of 60 covariates that could be used for propensity score matching (i.e. categorical or numerical data).

Many of these 60 covariates had a considerable amount of missing data. Data was assumed to be missing completely at random. Variables that were recorded as missing in over 50% of individuals were omitted from the analysis. These variables included house type (permanent/improvised), roof, floor, and wall material, number of people and families in the home, number of bedrooms and bathrooms, variables relating to employment status, expenditure on rent and transport, and receipt of pension, unemployment benefit, and alimony. It is conceivable that rent and transport expenditure could be important confounders of cure rate given the potential of cash transfers for mitigating catastrophic costs, but neither are conditionally associated with both outcome and exposure in the observed data and expenditure is represented by other retained variables [15].

The omission of variables with this level of missing data resulted in 45 covariates to be considered for use in propensity score estimation. A sensitivity analysis was run omitting all variables with over 25% missing data, which further omitted water expenditure and years of formal education. At both missing data thresholds, at least one proxy covariate remained under each node of the DAG such that no high-level construct was unrepresented by the available covariates.

### Propensity Score Matching

Without applying propensity score approaches or other approaches to control for confounding, it is likely that the values of the available covariates between the exposed and the unexposed (and those who experience or do not experience the outcome) vary, which potentially biases comparisons between groups. We wish to achieve a ‘balance’ in these values, similar to the balance produced by conventional randomisation procedures. We wish to first determine the likelihood of receiving BFP given the covariate values, which is represented by the propensity score. If the propensity score is then balanced between groups by matching, it is as though the covariates that were used to estimate the propensity score were themselves balanced [16].

Propensity scores were estimated by logistic regression. One of two criteria must be met for a variable to be included in this logistic regression: a) conditional association with the outcome given exposure or b) both association with exposure and conditional association with outcome given exposure [17]. These criteria apply to both mediators and confounders and can be determined from the DAG (Figure 1). All DAG nodes meet these criteria but housing and thus the covariates used to model the propensity score were all non-housing covariates meeting the missing data threshold. Quadratic forms of the continuous covariates were used in the logistic regression but sensitivity analyses were performed without including them. Two-way interactions between gender and all variables and age and all variables were also used, given it is likely that these covariates would differ in effect across strata.

Each patient who did not receive BFP was matched to a patient who did receive it closest in propensity score, within a particular ‘caliper’ of 0·1 standard deviations from the mean propensity score. Matching was done with replacement and multiple matches to minimise both bias and variance, following Caliendo & Kopeinig (2008) [18]. Multiple matches were weighted to form one matched control for each patient. Standardised mean differences and overlap plots were examined to assess whether balance was improved by matching.

Throughout the literature, complete cases are used for propensity score matching, and this is the approach used in this paper [18]. This reduced the dataset to 2167 individuals at the 50% missing data threshold and 3048 individuals at the 25% threshold.

### Estimating the Impact of BFP

Taking the difference of the proportion of cures between matched groups resulted in an estimate of the average effect of treatment on the treated (ATT), or the (causal) risk difference in the exposed. The procedure used in Abadie & Imbens (2011) was used to estimate the standard error of the ATT and thus the confidence intervals. The confidence intervals thus account for the uncertainty due to the matching procedure, but do not account for the uncertainty due to the fact that the estimated propensity score is itself a function of the data; this latter feature leads to conservative inferences.[19] The ATT was also estimated by a multiple imputation based sensitivity analysis, and point estimates from this are provided for comparative purposes in Appendix 2.

### Statistical Software

All analyses were conducted in R v3.4.1 and the MatchIt package was used for the propensity score matching procedure.

## Results

### Propensity Score Matching – Covariate Balance

A complete balance table is presented in Table 2 in Appendix 1 for the match produced by Model A for all covariates included in the propensity score matching exercise. There is good similarity of the covariates after matching, suggesting a reasonable balance was obtained between groups. Prior to matching, there were some imbalances found between BFP recipients and non-recipients on important covariates. Figure 2 presents the changes in standardised mean difference between those receiving BFP and those not receiving BFP before and after matching. Figure 3 presents overlap plots to demonstrate the similarity of the propensity score values between groups.

**Fig 2.**
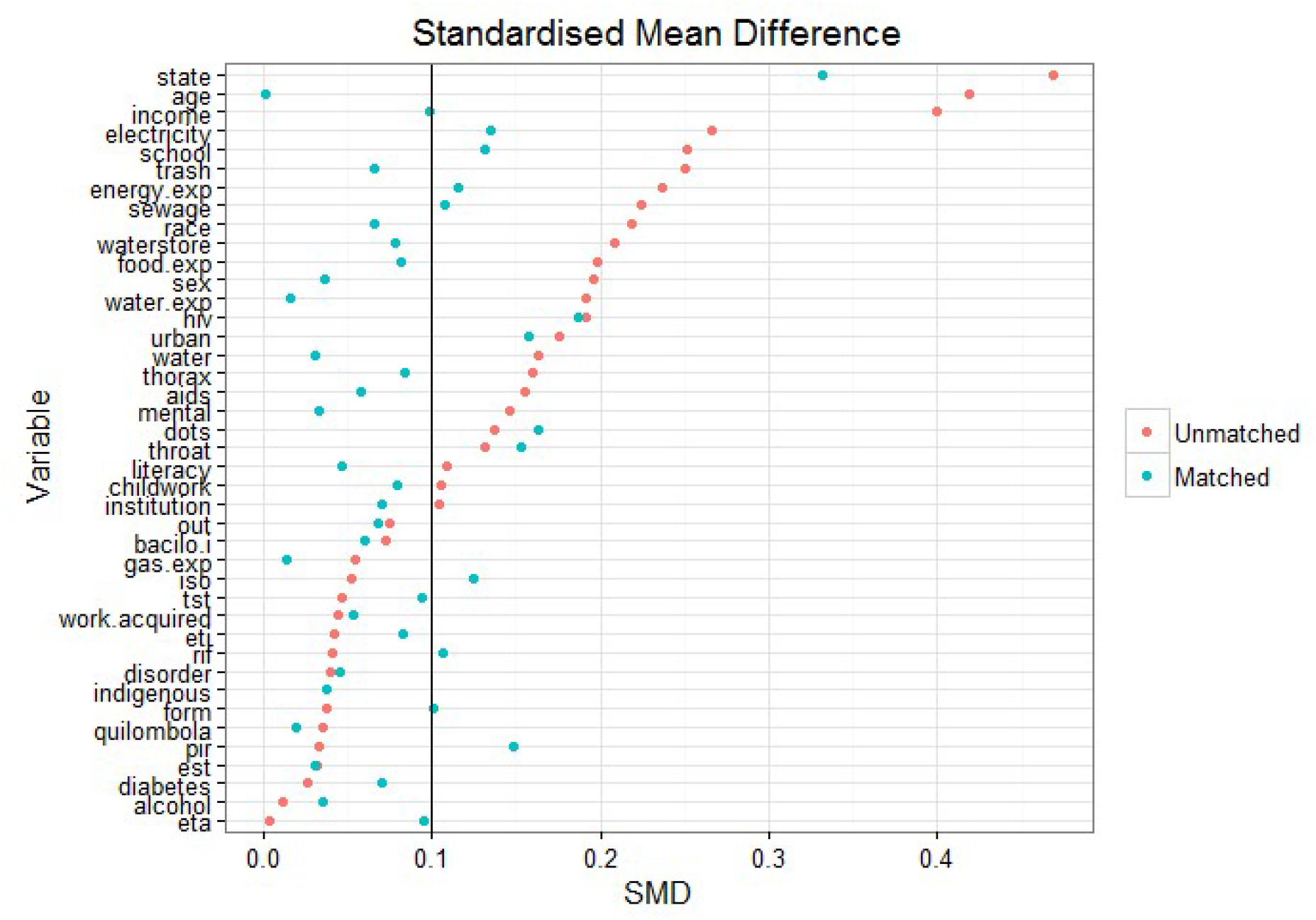
Standardised Mean Difference. The change in standardised mean difference in the matched and unmatched groups for each variable. A smaller difference indicates improved balance between groups; being below the threshold of 0.1 is conservatively considered to be effectively balanced. Balance has been largely improved by matching though some imbalance remains between groups.

**Fig 3.**
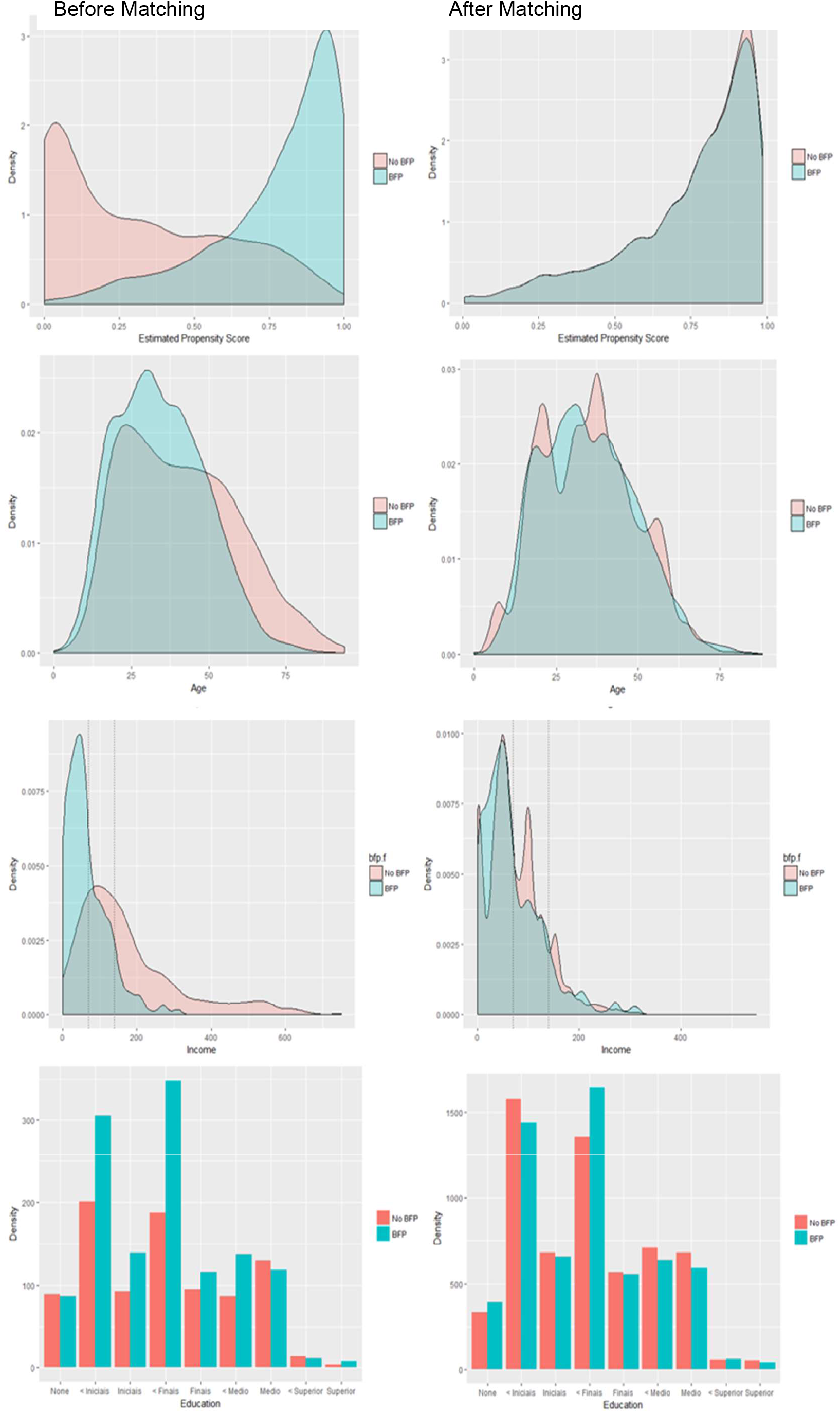
Overlap before and after matching. Overlap in estimated propensity scores between those receiving and those not receiving BFP before matching (top left) and after matching (top right). Overlap has been substantially improved by matching to treated (exposed) patients, suggestive of the groups being balanced on the propensity score. The region of overlap extends between 0 and 1. Also presented are similar plots of variable distribution before and after matching for income, age, and schooling (from top to bottom). Dotted lines on the income distributions mark the thresholds for BFP eligibility.

Propensity score matching in general resulted in improved balance of the values of covariates between cases and controls. A standardised mean difference of below 0.1 implies that groups do not differ greatly between values of the covariate [17]. Though the matching process only brought 50% of the imbalanced variables below this threshold, a large improvement was seen on the balance of important upstream covariates like age (0·42 to 0·01), income (0·40 to 0·09), and schooling (0·24 to 0·12). The change in distributions of these variables after matching can be seen in Figure 3. On average, those receiving BFP in the unmatched cohort were younger (34·5 vs. 41·3 years), poorer (65·2 vs. 197·4 reais per month), and less educated (89·2% vs 83.5% not completed secondary school).

From Figure 3, approximately 20·9% of TB patients fall under the 70 reais income threshold for unconditional receipt of BFP and therefore are theoretically eligible for the programme, but yet excluded from it. A further 29·4% fall under the 140 reais income threshold and could therefore potentially be eligible for BFP.

### Estimating the Impact of BFP

In total, four estimates of the ATT were produced (Table 1). Model A is the primary model of interest as it is the most complex model specification. Models B-D represent sensitivity analyses on Model A to investigate how sensitive the results are to simplifying changes to these modelling and missing data decisions.

**Table 1.**
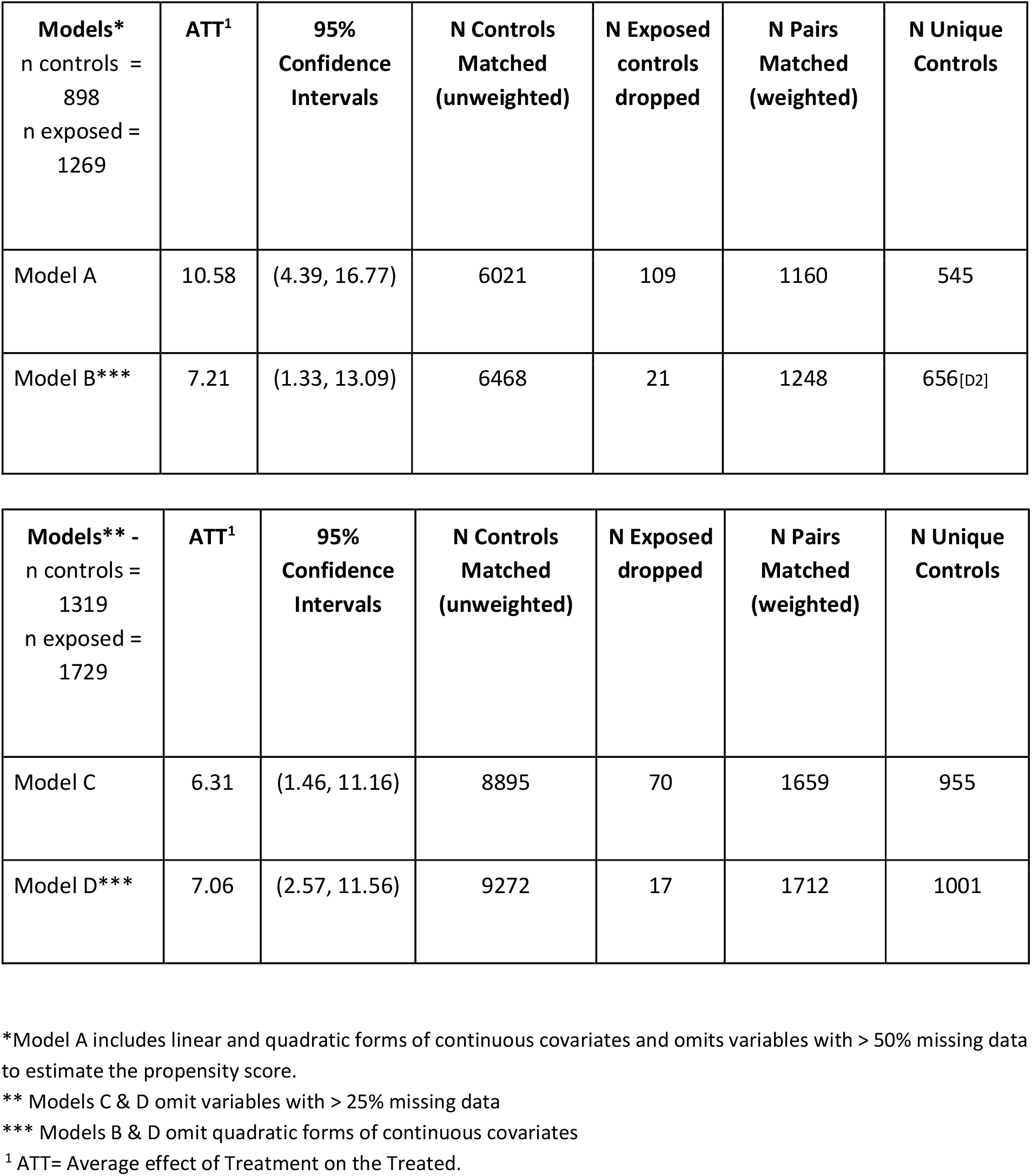
Results of propensity score matching estimates of the ATT for four models.

| **Models\***<br>n controls =<br>898<br>n exposed =<br>1269 | **ATT<sup>1</sup>** | **95%**<br>**Confidence**<br>**Intervals** | **N Controls**<br>**Matched**<br>**(unweighted)** | **N Exposed**<br>**controls**<br>**dropped** | **N Pairs**<br>**Matched**<br>**(weighted)** | **N Unique**<br>**Controls** |
| --- | --- | --- | --- | --- | --- | --- |
| Model A | 10.58 | (4.39, 16.77) | 6021 | 109 | 1160 | 545 |
| Model B*** | 7.21 | (1.33, 13.09) | 6468 | 21 | 1248 | 656[D2] |

| **Models\*\* -**<br>n controls =<br>1319<br>n exposed =<br>1729 | **ATT<sup>1</sup>** | **95%**<br>**Confidence**<br>**Intervals** | **N Controls**<br>**Matched**<br>**(unweighted)** | **N Exposed**<br>**dropped** | **N Pairs**<br>**Matched**<br>**(weighted)** | **N Unique**<br>**Controls** |
| --- | --- | --- | --- | --- | --- | --- |
| Model C | 6.31 | (1.46, 11.16) | 8895 | 70 | 1659 | 955 |
| Model D*** | 7.06 | (2.57, 11.56) | 9272 | 17 | 1712 | 1001 |

\*Model A includes linear and quadratic forms of continuous covariates and omits variables with > 50% missing data to estimate the propensity score.
\*\* Models C & D omit variables with > 25% missing data
\*\*\* Models B & D omit quadratic forms of continuous covariates
<sup>1</sup> ATT= Average effect of Treatment on the Treated.

The average effect of treatment on the treated from Model A was estimated to be 10·58 (95% CIs: 4·39, 16·77) (Table 1). Thus, amongst TB patients who receive BFP, we expect a cure rate 10·58% higher than if those patients had not received the benefit. The proportion cured in those who did not receive BFP was 76·6% compared to 87·2% in the BFP recipients. This average treatment effect is protective even when a simpler model is used and when the missing data threshold at which covariates are omitted is reduced to 25%, with ATT estimates between 6·31 and 7·21 (Table 1). It is also in broad agreement with an ATT point estimate of 7·22 obtained from a multiple imputation approach (Appendix 2).

## Discussion

### Summary – Interpretation of Results

This is the first study that uses a quasi-experimental approach to estimate the impact of a conditional cash transfer programme on TB cure rates. Across all models, results have shown a substantial absolute increase in TB cure rate (between 7-11%) amongst those who receive BFP. This suggests a consistent positive impact of BFP on a key indicator of TB control: cure rate. This is in line with Torrens et al. (2016), Durovni et al. (2017) and a few other previous studies evaluating the association between social protection and TB outcomes undertaken using less rigorous methodologies, which also demonstrate a protective effect of similar scale [7,20,21]. Similar propensity score approaches have already been used to evaluate the effect of cash transfers in HIV/AIDS, but not on TB [22].

Another important and somewhat unexpected finding of our analysis is that the profile of TB patients enrolled in BFP was not overtly dissimilar from TB patients that have not received BFP even before matching. Figure 2 suggests that the most imbalanced covariates for receipt of BFP (based on the standardised mean difference) were state of residence, income, age, and schooling. There may also be differences between recipients and non-recipients based on measures of the infrastructure of the local area (sewage, electricity, trash disposal). TB patients not benefiting from BFP transfers appear to be broadly similar to TB patients who are BFP recipients under a number of other sociodemographic characteristics, particularly on comorbidities such as diabetes and alcohol abuse, as well as on DOT prevalence (Table 2 in Appendix 1). This suggests there may be some shared vulnerability amongst TB patients but a disparity in access to BFP, likely heavily influenced by income and state. Our results show that up to 51·3% of patients may be theoretically eligible for BFP – this suggests that the income threshold for BFP is insufficiently specific to ensure access to vulnerable TB patients.

### Strengths & Limitations

The utilisation of quasi-experimental approach is a major strength of this paper. Quasi-experimental approaches like propensity score matching require fewer assumptions about the data than traditional parametric counterparts. The specification of the estimand and population parameters of interest are an additional strength to using propensity score matching, and the risk of bias from residual confounding is minimised compared to prior work by careful use of a DAG [23]. While the use of propensity scores for matching has recently drawn some criticism, the diagnostic plots demonstrated in Figures 2 & 3 show that issues of remaining imbalance are largely mitigated. Propensity score matching was also chosen as an alternative to regression adjustment or inverse probability of treatment weighting because of the framing of the study as a response to other work that did not compare to appropriate controls.

Indeed, a clear strength of this work is the comparability of the control group. As demonstrated in Figure 3, those in the exposed group and those in the control group have a very similar distribution of propensity to receive BFP. This overlap suggests that we are only comparing patients with similar covariate profiles: while some of the control patients may not be eligible on paper for Bolsa Familia, in the complex context of real-world receipt of BFP, the controls resemble almost exactly those patients who receive it and are representative of a broad range of TB patients from across Brazil. This is a methodological improvement over the control groups seen in prior work which greatly strengthens the quality of evidence available to policymakers [7,10].

The control group in Durovni et al. (2017) was taken from a pool of all TB patients rather than those who are registered in CadÚnico, and therefore some patients ineligible for BFP were included in the control group. The control group in Torrens et al. (2016) was taken from TB patients who were eligible in theory for BFP, but who had not received any money from the programme until after treatment. This control group had different characteristics to those TB patients not eligible for the programme on demographic and socioeconomic variables examined by the authors. Both of these control groups may have potentially biased the resulting estimate of proportion of patients cured attributable to BFP.

This quasi-experimental approach also implies the possibility of drawing causal conclusions. The estimand used in this study, the average treatment effect on the treated, can be given a causal interpretation if particular ‘identifying’ assumptions hold. The validity of these assumptions are briefly discussed in Appendix 3.

The major limitation to this work is the data quality. The missing data results in a relatively small sample size used for matching and we cannot rule out the possibility of residual confounding from covariates that are mostly missing or remain unbalanced. Remaining imbalance on the state variable suggests data may be missing conditionally at random on the state variable. As information on it is housed within a separate register, we were unable to assess the impact of the Family Health Strategy, though previous work suggests the effect of Bolsa Familia is independent of FHS coverage. While an approach combining multiple imputation and propensity score matching would have mitigated this problem, there remain many gaps in the literature on the practical implementation of these techniques together (see Appendix 2). Furthermore, the data linkage is cross-sectional and thus time-varying confounding cannot be accounted for with these data; better data availability longitudinally would allow for measurement on more direct measures of TB control, such as incidence.

The choice of a dichotomous outcome variable may be another limitation: non-cure outcomes include continued disease post regimen completion, treatment abandonment, death from TB, death from other causes, and development of MDR-TB, which may have heterogeneous risk factors. Loss to follow up and transferred cases are also not considered by this analysis – the analysis is agnostic about whether these patients were cured or not cured. The results may be different if each non-cure outcome were addressed in turn, but this would require a larger sample size and may be best addressed in a descriptive study.

### Policy Implications

These findings preliminarily suggest that: 1) there is a considerable proportion of TB patients eligible for BFP that for unknown reasons seem to be left out from the programme; 2) almost half of the TB patients will not be eligible for BFP according to income thresholds, and thus there is room for a more comprehensive or multidimensional targeting approach not only using income as eligibility criteria. Given the 7-11% absolute increase in cure rate seen amongst those receiving BFP from our work, from a global health rights perspective, it must be considered how best to deliver a protective programme to vulnerable patients.

Bolsa Familia was not designed to address specific diseases, not least TB, TB care or TB control objectives. Access could be expanded to TB patients by making TB status an eligibility criterion for the programme or by making efforts to make BFP more TB-sensitive by using a more inclusive, albeit non-stigmatising, targeting strategy. According to our findings, better BFP coverage and thus improved TB outcomes among TB patients could be achieved by simply facilitating access to BFP benefits from patients that are already eligible by definition for the programme, but not actually receiving the programme benefits. To this purpose, further research is urgently needed to understand determinants of access to BFP, especially for TB patients. Understanding the likely impact on the Brazilian TB epidemic if these barriers to programme receipt at the individual and system level were removed will be necessary.

It is likely that the removal of these barriers may require the implementation of more efficient BFP delivery models, including the ‘single window’ approach which entails an integrated delivery of TB care services and social protection. According to this model, the access to the most appropriate social protection schemes is determined and facilitated at the primary health care level where ad hoc staff (e.g. social workers) are trained to assess the social protection needs of TB patients and provide information, legal and administrative advices, and referrals to various services so to allow patients to access benefits from one ‘single window’ without having to navigate across complex and multiple service points [24].

Another emerging model for the delivery of social protection is the ‘cash plus’ model in which the provision of cash transfers is combined with another form of social support when the provision of in kind benefits is not deemed sufficient to reduce households’ vulnerabilities (including health related vulnerabilities) [25].

In the case of TB in Brazil, this ‘plus’ component can be represented by a top up of the cash benefit to account for the TB-related catastrophic costs incurred by the households; or the provision of a food basket to improve nutrition of cash beneficiaries and therefore their treatment outcome; or the improvement of housing and ventilation conditions to interrupt intra-household transmission of TB. To identify the most relevant ‘intensifier’ of any cash transfer intervention it will be essential also to understand thoroughly the most likely pathway through which this impact takes place. This requires the development of a setting-specific, epidemiologically driven conceptual framework and a more comprehensive collection of data for the variables in the causal pathway.

### Conclusions

Overall, the strength of evidence and size of effect of the ATT estimated in this work suggest that a paradigm shift is needed towards viewing expansion of social protection to a wider population of TB patients as a valid mechanism for improving TB outcomes beyond the traditional biomedical approach. This must be part of a broader push for changing practice in TB control towards working within existing development models and infrastructures. It is essential that, like in this work, recent developments in quasi-experimental methodology continue to be integrated with the evidence base for bold policies in development. With stronger evidence available, the rapid implementation of bold policies may be justified in TB contexts and the global public health community will be a large step closer to achieving the aims of the WHO’s End-TB Strategy.

Control patients were many-to-one matched to treatment patients with replacement (Controls matched unweighted), excluding those who were dropped by the caliper (Exposed controls dropped). Matched controls were weighted to form one matched comparator for each treatment patient. These matched comparator patients were matched to the treatment patients to form matched pairs (Pairs of controls and treated cases matched). The number of pairs may thus be higher than the total initial sample size as some controls were used more than once and some were not used at all (Number of unique controls).

## Appendix 1 - Table 2. Baseline Characteristics

The following table presents the baseline characteristics of the unmatched population and the same characteristics after matching. Variables correspond to the nodes of the DAG in Figure 1.

**Table 2:**
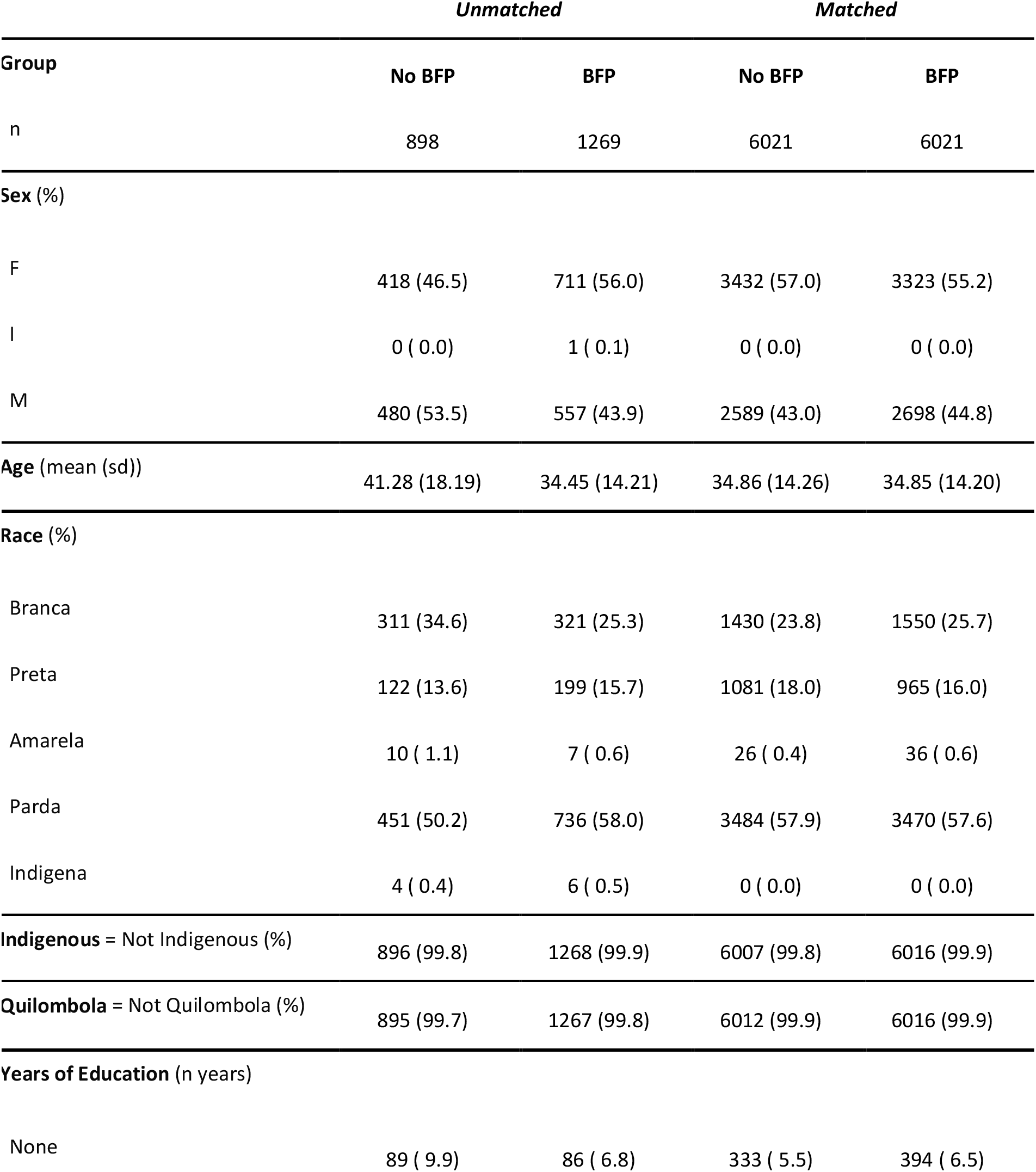
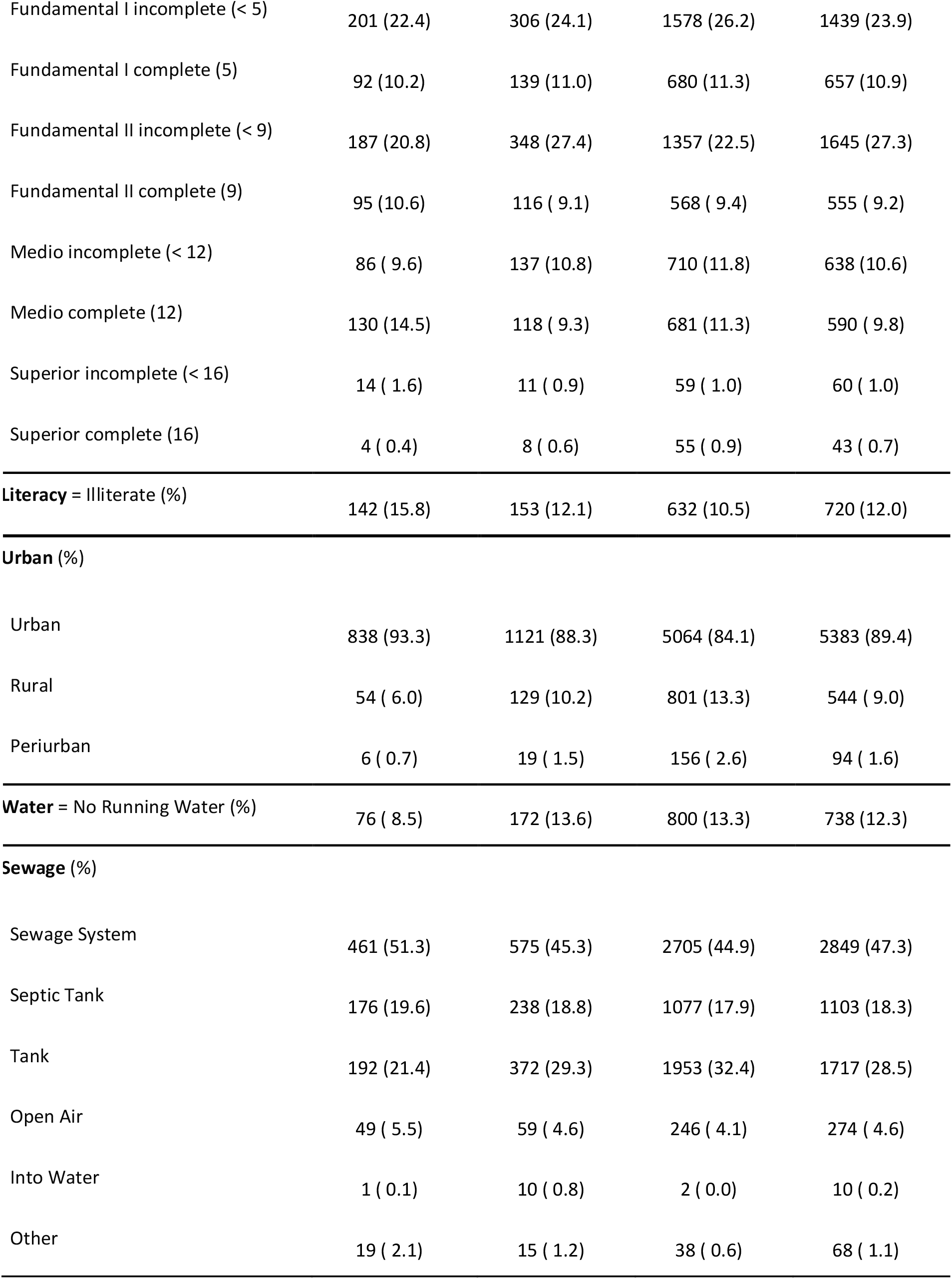
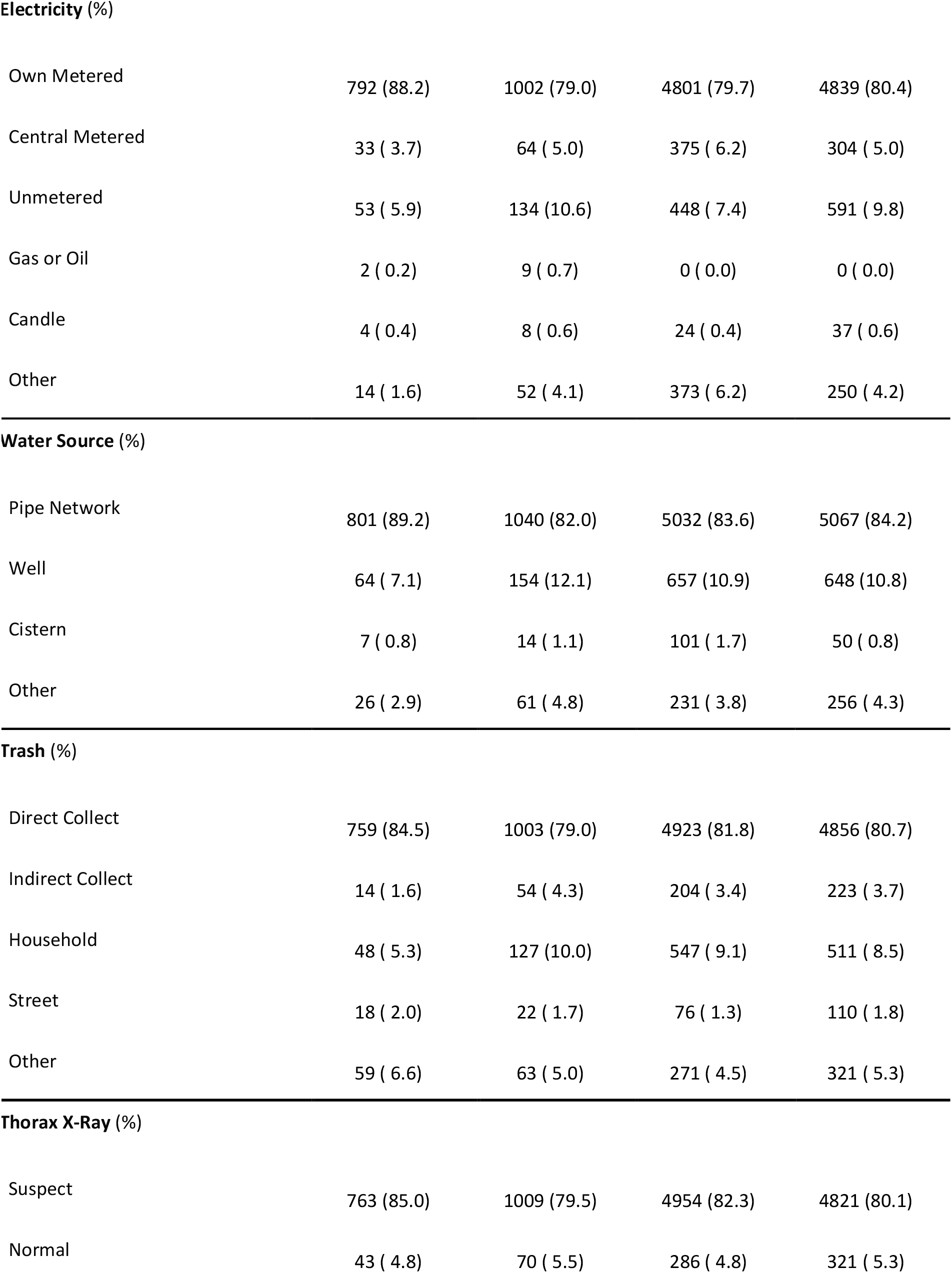
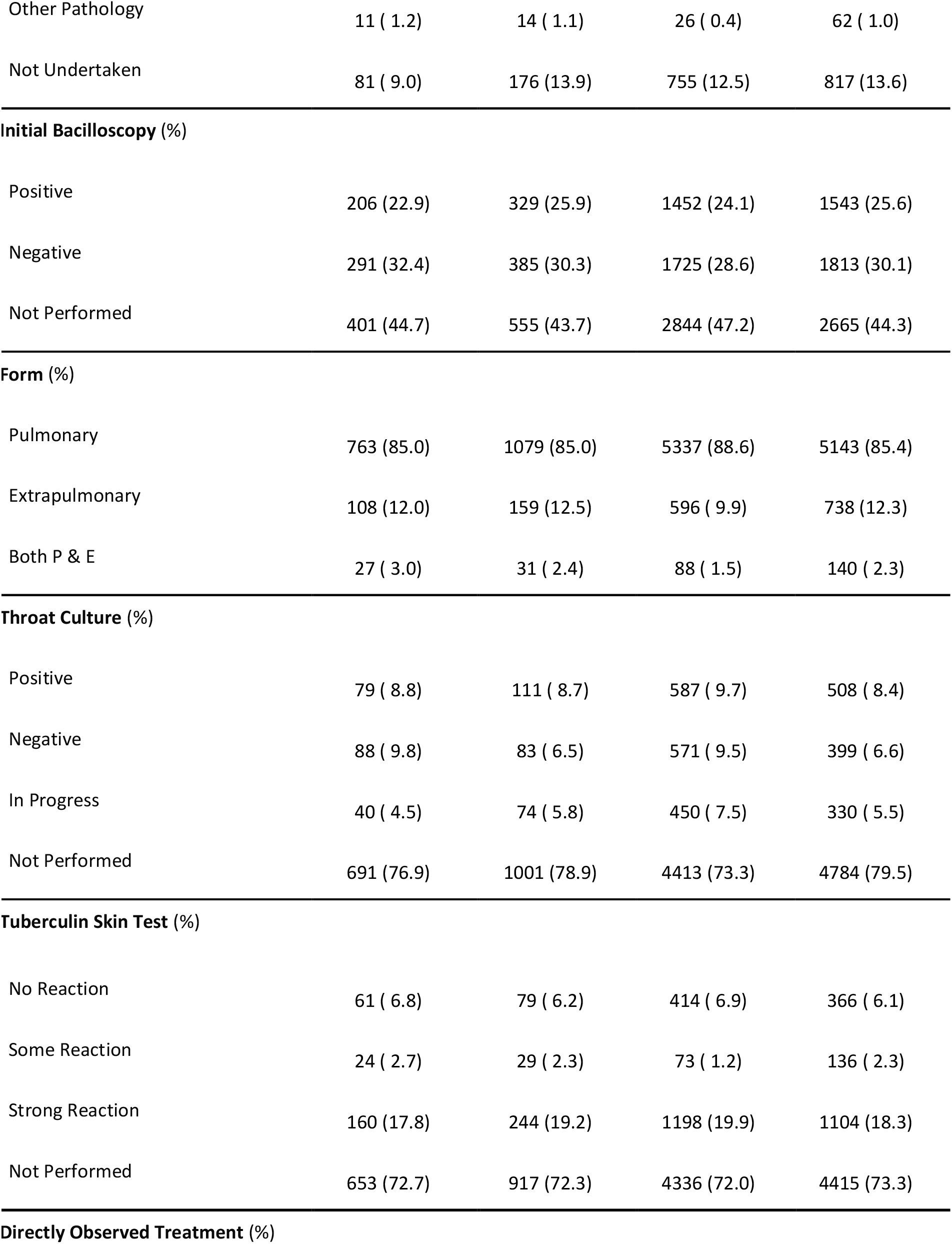
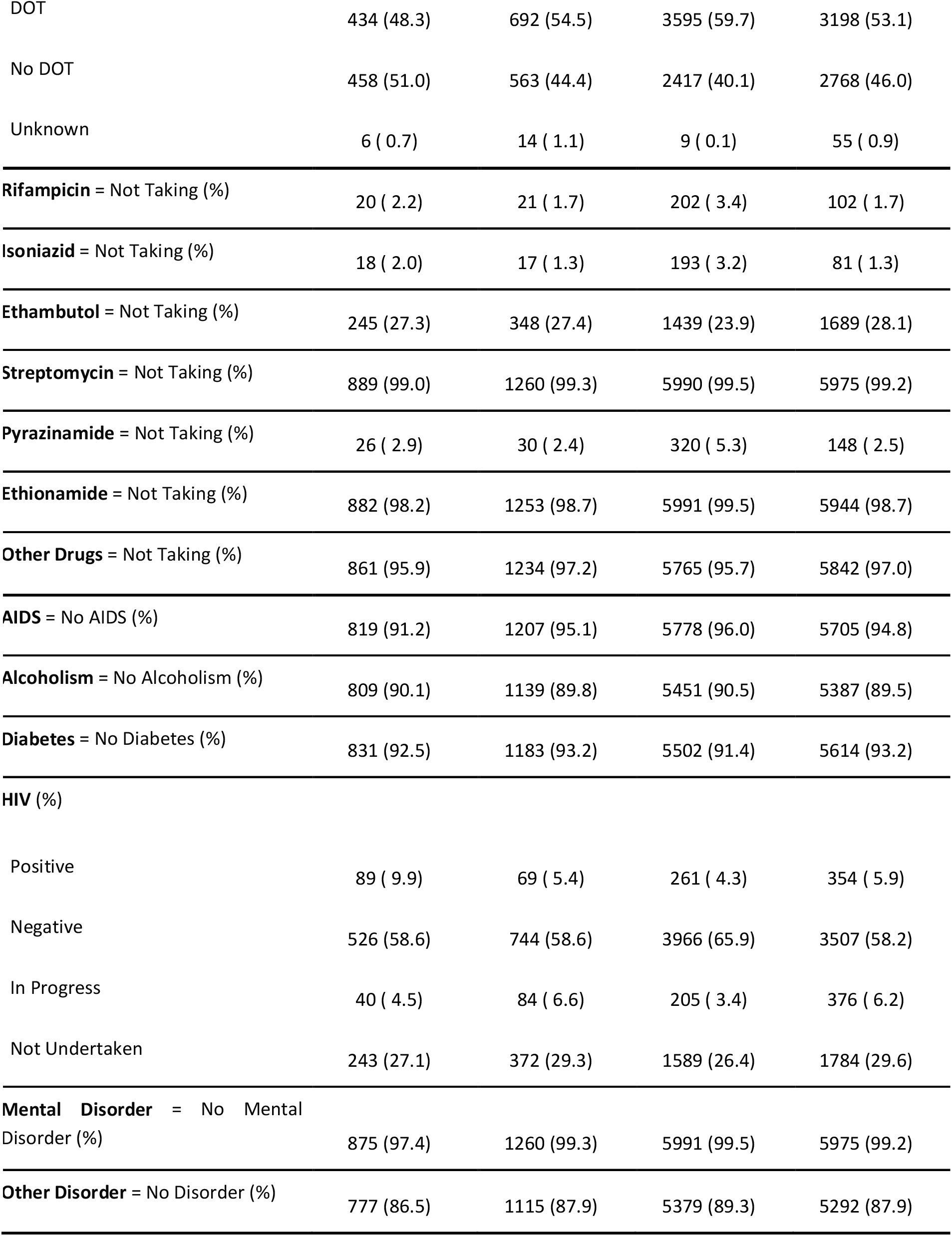
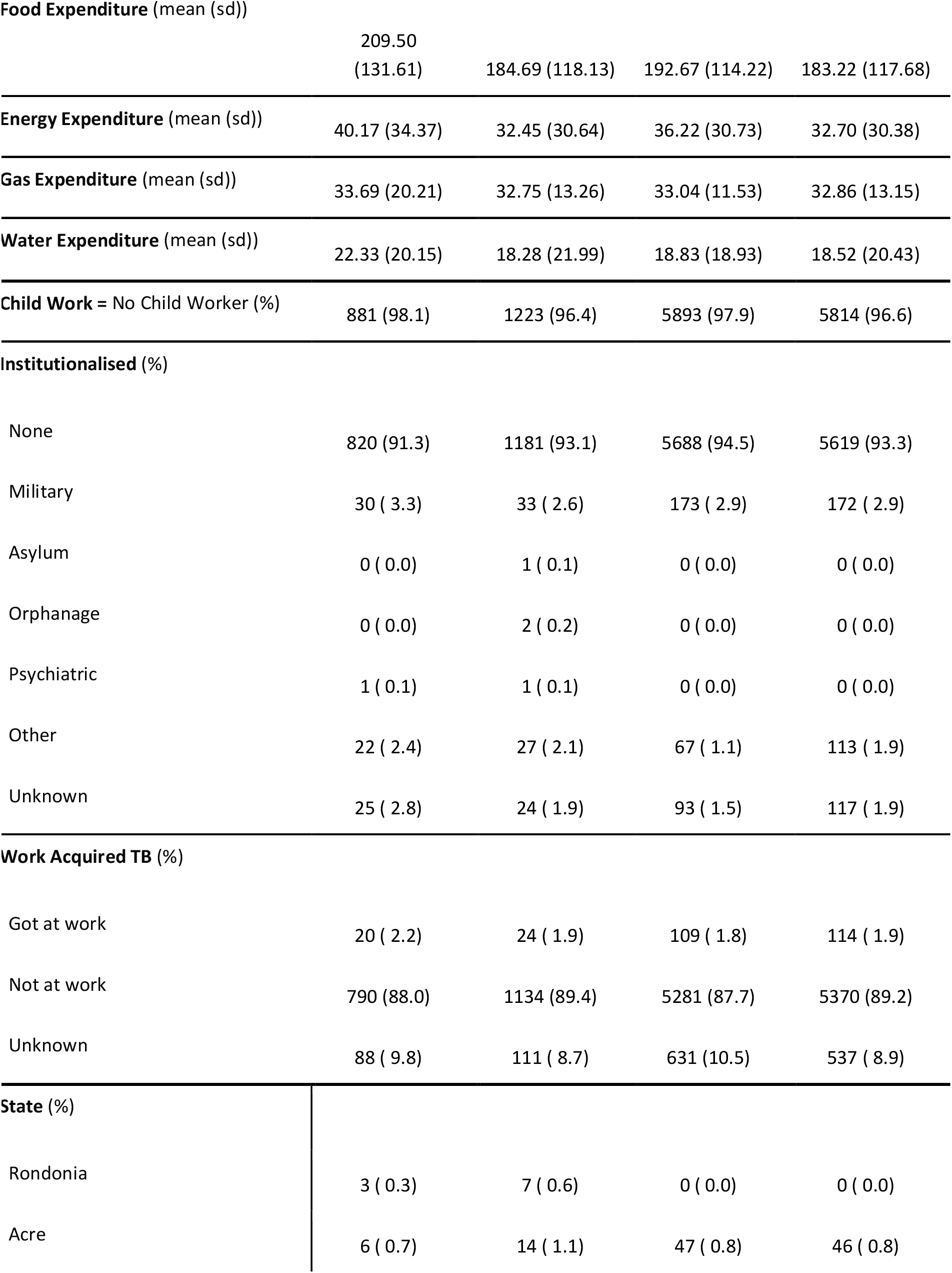
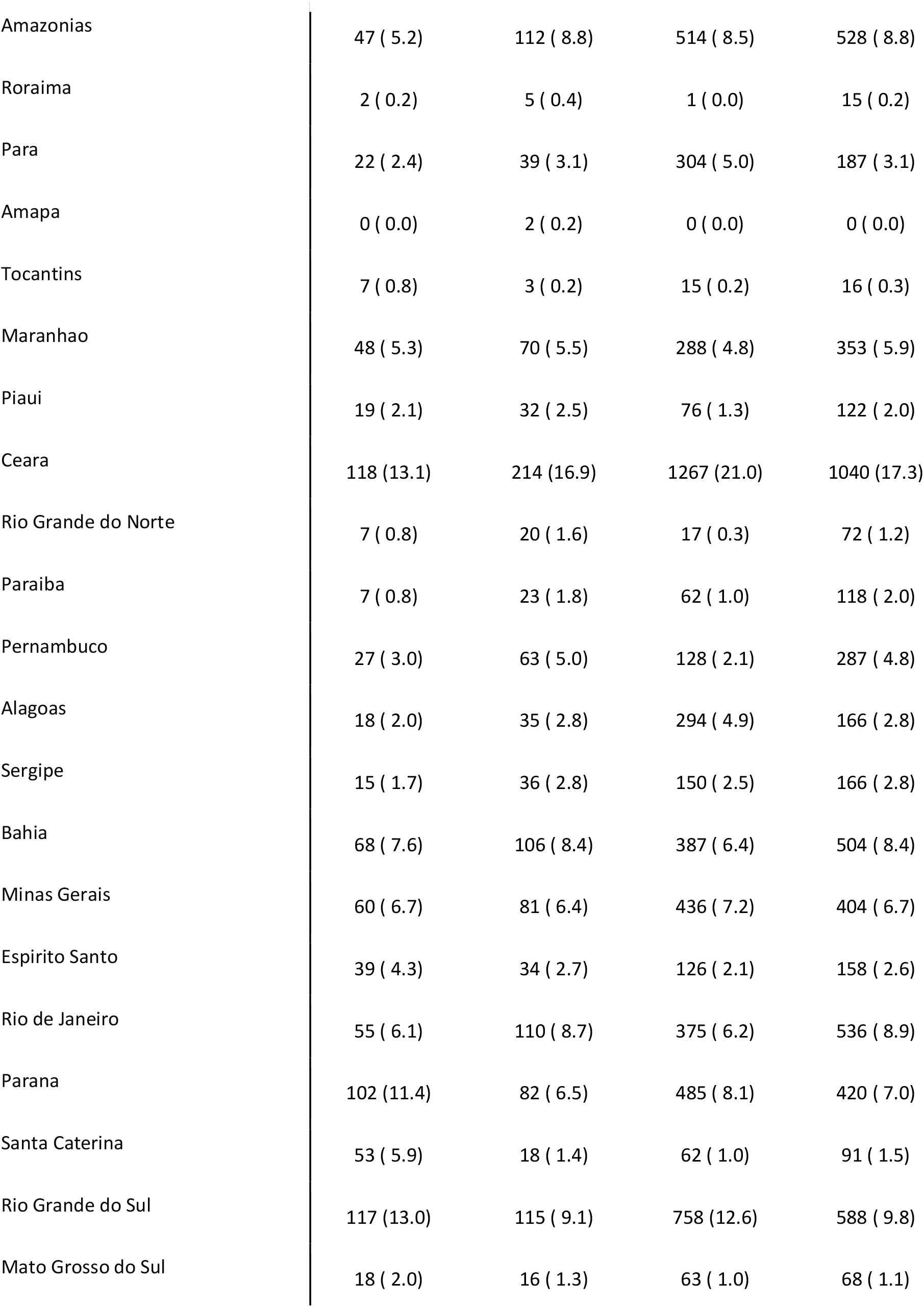
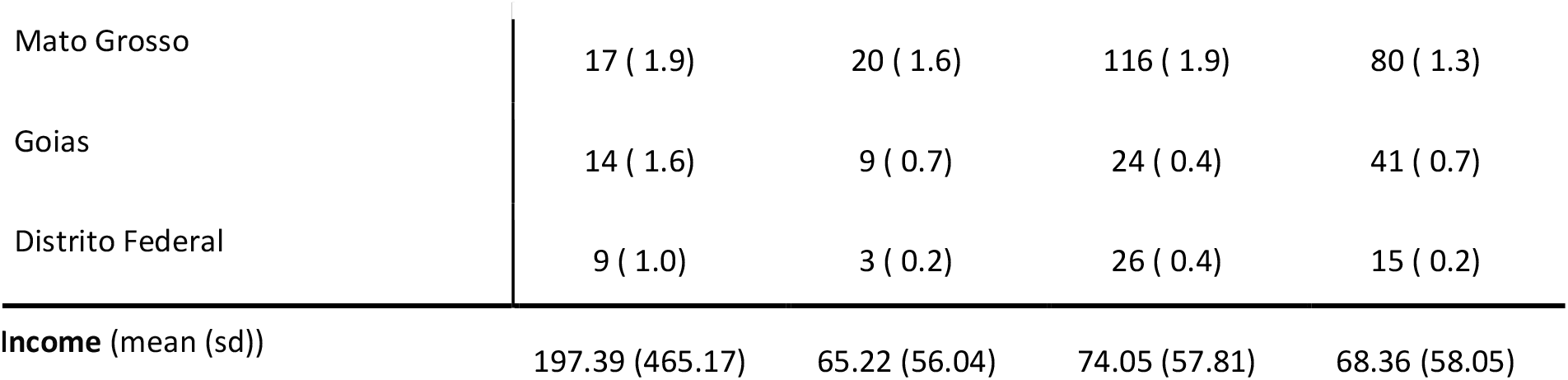
Table of balance on covariates both before and after matching, stratified by receipt of BFP. Covariates are grouped by DAG node including *Sex & Age* (Sex, Age), *Race* (Race, Indigenous, Quilombola), *Education* (Education Level, Literacy), *Local Area* (Urban, Running Water, Sewage, Electricity, Water Store, Trash), *Type of TB* (Thorax X-Ray, Initial Bacilloscopy, Form, Throat Culture, Tuberculin Skin Test), *Directly Observed Treatment* (DOT), *Drugs* (Rifampicin, Isoniazid, Ethambutol, Streptomycin, Pyrazinamide, Ethionamide, Other Drugs), *Comorbidities* (AIDS, Alcoholism, Diabetes, HIV, Mental Disorder, Other Disorder), *Expenditure* (on Food, Energy, Gas, and Water), *Social Vulnerability* (Child Worker, Institutionalised, Work Acquired TB), *State*, and *Income*. Where duplicate variables existed, SINAN was used preferentially.

| Group | *Unmatched* |  | *Matched* |  |
| --- | --- | --- | --- | --- |
|  | No BFP | BFP | No BFP | BFP |
| n | 898 | 1269 | 6021 | 6021 |
| **Sex (%)** |  |  |  |  |
| F | 418 (46.5) | 711 (56.0) | 3432 (57.0) | 3323 (55.2) |
| I | 0 ( 0.0) | 1 ( 0.1) | 0 ( 0.0) | 0 ( 0.0) |
| M | 480 (53.5) | 557 (43.9) | 2589 (43.0) | 2698 (44.8) |
| **Age (mean (sd))** | 41.28 (18.19) | 34.45 (14.21) | 34.86 (14.26) | 34.85 (14.20) |
| **Race (%)** |  |  |  |  |
| Branca | 311 (34.6) | 321 (25.3) | 1430 (23.8) | 1550 (25.7) |
| Preta | 122 (13.6) | 199 (15.7) | 1081 (18.0) | 965 (16.0) |
| Amarela | 10 ( 1.1) | 7 ( 0.6) | 26 ( 0.4) | 36 ( 0.6) |
| Parda | 451 (50.2) | 736 (58.0) | 3484 (57.9) | 3470 (57.6) |
| Indigena | 4 ( 0.4) | 6 ( 0.5) | 0 ( 0.0) | 0 ( 0.0) |
| **Indigenous = Not Indigenous (%)** | 896 (99.8) | 1268 (99.9) | 6007 (99.8) | 6016 (99.9) |
| **Quilombola = Not Quilombola (%)** | 895 (99.7) | 1267 (99.8) | 6012 (99.9) | 6016 (99.9) |
| **Years of Education (n years)** |  |  |  |  |
| None | 89 ( 9.9) | 86 ( 6.8) | 333 ( 5.5) | 394 ( 6.5) |
| Fundamental I incomplete (< 5) | 201 (22.4) | 306 (24.1) | 1578 (26.2) | 1439 (23.9) |
| Fundamental I complete (5) | 92 (10.2) | 139 (11.0) | 680 (11.3) | 657 (10.9) |
| Fundamental II incomplete (< 9) | 187 (20.8) | 348 (27.4) | 1357 (22.5) | 1645 (27.3) |
| Fundamental II complete (9) | 95 (10.6) | 116 ( 9.1) | 568 ( 9.4) | 555 ( 9.2) |
| Medio incomplete (< 12) | 86 ( 9.6) | 137 (10.8) | 710 (11.8) | 638 (10.6) |
| Medio complete (12) | 130 (14.5) | 118 ( 9.3) | 681 (11.3) | 590 ( 9.8) |
| Superior incomplete (< 16) | 14 ( 1.6) | 11 ( 0.9) | 59 ( 1.0) | 60 ( 1.0) |
| Superior complete (16) | 4 ( 0.4) | 8 ( 0.6) | 55 ( 0.9) | 43 ( 0.7) |
| **Literacy = Illiterate (%)** | **142 (15.8)** | **153 (12.1)** | **632 (10.5)** | **720 (12.0)** |
| **Urban (%)** |  |  |  |  |
| Urban | 838 (93.3) | 1121 (88.3) | 5064 (84.1) | 5383 (89.4) |
| Rural | 54 ( 6.0) | 129 (10.2) | 801 (13.3) | 544 ( 9.0) |
| Periurban | 6 ( 0.7) | 19 ( 1.5) | 156 ( 2.6) | 94 ( 1.6) |
| **Water = No Running Water (%)** | **76 ( 8.5)** | **172 (13.6)** | **800 (13.3)** | **738 (12.3)** |
| **Sewage (%)** |  |  |  |  |
| Sewage System | 461 (51.3) | 575 (45.3) | 2705 (44.9) | 2849 (47.3) |
| Septic Tank | 176 (19.6) | 238 (18.8) | 1077 (17.9) | 1103 (18.3) |
| Tank | 192 (21.4) | 372 (29.3) | 1953 (32.4) | 1717 (28.5) |
| Open Air | 49 ( 5.5) | 59 ( 4.6) | 246 ( 4.1) | 274 ( 4.6) |
| Into Water | 1 ( 0.1) | 10 ( 0.8) | 2 ( 0.0) | 10 ( 0.2) |
| Other | 19 ( 2.1) | 15 ( 1.2) | 38 ( 0.6) | 68 ( 1.1) |
| **Electricity (%)** |  |  |  |  |
| Own Metered | 792 (88.2) | 1002 (79.0) | 4801 (79.7) | 4839 (80.4) |
| Central Metered | 33 ( 3.7) | 64 ( 5.0) | 375 ( 6.2) | 304 ( 5.0) |
| Unmetered | 53 ( 5.9) | 134 (10.6) | 448 ( 7.4) | 591 ( 9.8) |
| Gas or Oil | 2 ( 0.2) | 9 ( 0.7) | 0 ( 0.0) | 0 ( 0.0) |
| Candle | 4 ( 0.4) | 8 ( 0.6) | 24 ( 0.4) | 37 ( 0.6) |
| Other | 14 ( 1.6) | 52 ( 4.1) | 373 ( 6.2) | 250 ( 4.2) |
| **Water Source (%)** |  |  |  |  |
| Pipe Network | 801 (89.2) | 1040 (82.0) | 5032 (83.6) | 5067 (84.2) |
| Well | 64 ( 7.1) | 154 (12.1) | 657 (10.9) | 648 (10.8) |
| Cistern | 7 ( 0.8) | 14 ( 1.1) | 101 ( 1.7) | 50 ( 0.8) |
| Other | 26 ( 2.9) | 61 ( 4.8) | 231 ( 3.8) | 256 ( 4.3) |
| **Trash (%)** |  |  |  |  |
| Direct Collect | 759 (84.5) | 1003 (79.0) | 4923 (81.8) | 4856 (80.7) |
| Indirect Collect | 14 ( 1.6) | 54 ( 4.3) | 204 ( 3.4) | 223 ( 3.7) |
| Household | 48 ( 5.3) | 127 (10.0) | 547 ( 9.1) | 511 ( 8.5) |
| Street | 18 ( 2.0) | 22 ( 1.7) | 76 ( 1.3) | 110 ( 1.8) |
| Other | 59 ( 6.6) | 63 ( 5.0) | 271 ( 4.5) | 321 ( 5.3) |
| **Thorax X-Ray (%)** |  |  |  |  |
| Suspect | 763 (85.0) | 1009 (79.5) | 4954 (82.3) | 4821 (80.1) |
| Normal | 43 ( 4.8) | 70 ( 5.5) | 286 ( 4.8) | 321 ( 5.3) |
| Other Pathology | 11 ( 1.2) | 14 ( 1.1) | 26 ( 0.4) | 62 ( 1.0) |
| Not Undertaken | 81 ( 9.0) | 176 (13.9) | 755 (12.5) | 817 (13.6) |
| **Initial Bacilloscopy (%)** |  |  |  |  |
| Positive | 206 (22.9) | 329 (25.9) | 1452 (24.1) | 1543 (25.6) |
| Negative | 291 (32.4) | 385 (30.3) | 1725 (28.6) | 1813 (30.1) |
| Not Performed | 401 (44.7) | 555 (43.7) | 2844 (47.2) | 2665 (44.3) |
| **Form (%)** |  |  |  |  |
| Pulmonary | 763 (85.0) | 1079 (85.0) | 5337 (88.6) | 5143 (85.4) |
| Extrapulmonary | 108 (12.0) | 159 (12.5) | 596 ( 9.9) | 738 (12.3) |
| Both P & E | 27 ( 3.0) | 31 ( 2.4) | 88 ( 1.5) | 140 ( 2.3) |
| **Throat Culture (%)** |  |  |  |  |
| Positive | 79 ( 8.8) | 111 ( 8.7) | 587 ( 9.7) | 508 ( 8.4) |
| Negative | 88 ( 9.8) | 83 ( 6.5) | 571 ( 9.5) | 399 ( 6.6) |
| In Progress | 40 ( 4.5) | 74 ( 5.8) | 450 ( 7.5) | 330 ( 5.5) |
| Not Performed | 691 (76.9) | 1001 (78.9) | 4413 (73.3) | 4784 (79.5) |
| **Tuberculin Skin Test (%)** |  |  |  |  |
| No Reaction | 61 ( 6.8) | 79 ( 6.2) | 414 ( 6.9) | 366 ( 6.1) |
| Some Reaction | 24 ( 2.7) | 29 ( 2.3) | 73 ( 1.2) | 136 ( 2.3) |
| Strong Reaction | 160 (17.8) | 244 (19.2) | 1198 (19.9) | 1104 (18.3) |
| Not Performed | 653 (72.7) | 917 (72.3) | 4336 (72.0) | 4415 (73.3) |
| **Directly Observed Treatment (%)** |  |  |  |  |
| DOT | 434 (48.3) | 692 (54.5) | 3595 (59.7) | 3198 (53.1) |
| No DOT | 458 (51.0) | 563 (44.4) | 2417 (40.1) | 2768 (46.0) |
| Unknown | 6 ( 0.7) | 14 ( 1.1) | 9 ( 0.1) | 55 ( 0.9) |
| **Rifampicin = Not Taking (%)** | 20 ( 2.2) | 21 ( 1.7) | 202 ( 3.4) | 102 ( 1.7) |
| **Isoniazid = Not Taking (%)** | 18 ( 2.0) | 17 ( 1.3) | 193 ( 3.2) | 81 ( 1.3) |
| **Ethambutol = Not Taking (%)** | 245 (27.3) | 348 (27.4) | 1439 (23.9) | 1689 (28.1) |
| **Streptomycin = Not Taking (%)** | 889 (99.0) | 1260 (99.3) | 5990 (99.5) | 5975 (99.2) |
| **Pyrazinamide = Not Taking (%)** | 26 ( 2.9) | 30 ( 2.4) | 320 ( 5.3) | 148 ( 2.5) |
| **Ethionamide = Not Taking (%)** | 882 (98.2) | 1253 (98.7) | 5991 (99.5) | 5944 (98.7) |
| **Other Drugs = Not Taking (%)** | 861 (95.9) | 1234 (97.2) | 5765 (95.7) | 5842 (97.0) |
| **AIDS = No AIDS (%)** | 819 (91.2) | 1207 (95.1) | 5778 (96.0) | 5705 (94.8) |
| **Alcoholism = No Alcoholism (%)** | 809 (90.1) | 1139 (89.8) | 5451 (90.5) | 5387 (89.5) |
| **Diabetes = No Diabetes (%)** | 831 (92.5) | 1183 (93.2) | 5502 (91.4) | 5614 (93.2) |
| **HIV (%)** |  |  |  |  |
| Positive | 89 ( 9.9) | 69 ( 5.4) | 261 ( 4.3) | 354 ( 5.9) |
| Negative | 526 (58.6) | 744 (58.6) | 3966 (65.9) | 3507 (58.2) |
| In Progress | 40 ( 4.5) | 84 ( 6.6) | 205 ( 3.4) | 376 ( 6.2) |
| Not Undertaken | 243 (27.1) | 372 (29.3) | 1589 (26.4) | 1784 (29.6) |
| **Mental Disorder = No Mental Disorder (%)** | 875 (97.4) | 1260 (99.3) | 5991 (99.5) | 5975 (99.2) |
| **Other Disorder = No Disorder (%)** | 777 (86.5) | 1115 (87.9) | 5379 (89.3) | 5292 (87.9) |

**Food Expenditure (mean (sd))**
|  |  |  |  |
| --- | --- | --- | --- |
| 209.50<br>(131.61) | 184.69 (118.13) | 192.67 (114.22) | 183.22 (117.68) |

**Energy Expenditure (mean (sd))**
|  |  |  |  |
| --- | --- | --- | --- |
| 40.17 (34.37) | 32.45 (30.64) | 36.22 (30.73) | 32.70 (30.38) |

**Gas Expenditure (mean (sd))**
|  |  |  |  |
| --- | --- | --- | --- |
| 33.69 (20.21) | 32.75 (13.26) | 33.04 (11.53) | 32.86 (13.15) |

**Water Expenditure (mean (sd))**
|  |  |  |  |
| --- | --- | --- | --- |
| 22.33 (20.15) | 18.28 (21.99) | 18.83 (18.93) | 18.52 (20.43) |

**Child Work = No Child Worker (%)**
|  |  |  |  |
| --- | --- | --- | --- |
| 881 (98.1) | 1223 (96.4) | 5893 (97.9) | 5814 (96.6) |
| **Institutionalised (%)** |  |  |  |  |
| None | 820 (91.3) | 1181 (93.1) | 5688 (94.5) | 5619 (93.3) |
| Military | 30 ( 3.3) | 33 ( 2.6) | 173 ( 2.9) | 172 ( 2.9) |
| Asylum | 0 ( 0.0) | 1 ( 0.1) | 0 ( 0.0) | 0 ( 0.0) |
| Orphanage | 0 ( 0.0) | 2 ( 0.2) | 0 ( 0.0) | 0 ( 0.0) |
| Psychiatric | 1 ( 0.1) | 1 ( 0.1) | 0 ( 0.0) | 0 ( 0.0) |
| Other | 22 ( 2.4) | 27 ( 2.1) | 67 ( 1.1) | 113 ( 1.9) |
| Unknown | 25 ( 2.8) | 24 ( 1.9) | 93 ( 1.5) | 117 ( 1.9) |
| **Work Acquired TB (%)** |  |  |  |  |
| Got at work | 20 ( 2.2) | 24 ( 1.9) | 109 ( 1.8) | 114 ( 1.9) |
| Not at work | 790 (88.0) | 1134 (89.4) | 5281 (87.7) | 5370 (89.2) |
| Unknown | 88 ( 9.8) | 111 ( 8.7) | 631 (10.5) | 537 ( 8.9) |

**Table 2:** Table of balance on covariates both before and after matching, stratified by receipt of BFP.
|  |  |  |  |  |
| --- | --- | --- | --- | --- |
| Rondonia | 3 ( 0.3) | 7 ( 0.6) | 0 ( 0.0) | 0 ( 0.0) |
| Acre | 6 ( 0.7) | 14 ( 1.1) | 47 ( 0.8) | 46 ( 0.8) |
| Amazonias | 47 ( 5.2) | 112 ( 8.8) | 514 ( 8.5) | 528 ( 8.8) |
| Roraima | 2 ( 0.2) | 5 ( 0.4) | 1 ( 0.0) | 15 ( 0.2) |
| Para | 22 ( 2.4) | 39 ( 3.1) | 304 ( 5.0) | 187 ( 3.1) |
| Amapa | 0 ( 0.0) | 2 ( 0.2) | 0 ( 0.0) | 0 ( 0.0) |
| Tocantins | 7 ( 0.8) | 3 ( 0.2) | 15 ( 0.2) | 16 ( 0.3) |
| Maranhao | 48 ( 5.3) | 70 ( 5.5) | 288 ( 4.8) | 353 ( 5.9) |
| Piaui | 19 ( 2.1) | 32 ( 2.5) | 76 ( 1.3) | 122 ( 2.0) |
| Ceara | 118 (13.1) | 214 (16.9) | 1267 (21.0) | 1040 (17.3) |
| Rio Grande do Norte | 7 ( 0.8) | 20 ( 1.6) | 17 ( 0.3) | 72 ( 1.2) |
| Paraiba | 7 ( 0.8) | 23 ( 1.8) | 62 ( 1.0) | 118 ( 2.0) |
| Pernambuco | 27 ( 3.0) | 63 ( 5.0) | 128 ( 2.1) | 287 ( 4.8) |
| Alagoas | 18 ( 2.0) | 35 ( 2.8) | 294 ( 4.9) | 166 ( 2.8) |
| Sergipe | 15 ( 1.7) | 36 ( 2.8) | 150 ( 2.5) | 166 ( 2.8) |
| Bahia | 68 ( 7.6) | 106 ( 8.4) | 387 ( 6.4) | 504 ( 8.4) |
| Minas Gerais | 60 ( 6.7) | 81 ( 6.4) | 436 ( 7.2) | 404 ( 6.7) |
| Espirito Santo | 39 ( 4.3) | 34 ( 2.7) | 126 ( 2.1) | 158 ( 2.6) |
| Rio de Janeiro | 55 ( 6.1) | 110 ( 8.7) | 375 ( 6.2) | 536 ( 8.9) |
| Parana | 102 (11.4) | 82 ( 6.5) | 485 ( 8.1) | 420 ( 7.0) |
| Santa Caterina | 53 ( 5.9) | 18 ( 1.4) | 62 ( 1.0) | 91 ( 1.5) |
| Rio Grande do Sul | 117 (13.0) | 115 ( 9.1) | 758 (12.6) | 588 ( 9.8) |
| Mato Grosso do Sul | 18 ( 2.0) | 16 ( 1.3) | 63 ( 1.0) | 68 ( 1.1) |
| Mato Grosso | 17 ( 1.9) | 20 ( 1.6) | 116 ( 1.9) | 80 ( 1.3) |
| Goiás | 14 ( 1.6) | 9 ( 0.7) | 24 ( 0.4) | 41 ( 0.7) |
| Distrito Federal | 9 ( 1.0) | 3 ( 0.2) | 26 ( 0.4) | 15 ( 0.2) |
| **Income** (mean (sd)) | 197.39 (465.17) | 65.22 (56.04) | 74.05 (57.81) | 68.36 (58.05) |

## Appendix 2 – Missing Data

Though it was not the primary analytical method for this work, a confirmatory sensitivity analysis using multiple imputation was undertaken using the MICE (multiple imputation by chained equations) approach, as implemented in the MICE package in R [26,27]. The MICE package defaults of predictive mean matching and polytomous regression were used as the imputation methods for numeric and categorical variables respectively, creating 5 multiply imputed datasets.

The literature is still unclear as to whether the pooling of propensity scores themselves or the pooling of treatment estimates is the better approach after multiple imputation. Here, we followed Leyrat et al. (2016) and applied Rubin’s rules to pool the ATT estimates from each imputed dataset [28,29]. The ATT estimates were based on a comparison between groups that were matched on the propensity score estimated by the same model specification used for Model A. The resulting estimated ATT was 7.22, in broad agreement with other results.

An approach combining multiple imputation and propensity score methods was not used for the primary analysis due to numerous unresolved questions that admit the possibility for an unknown amount of bias with regards to the estimation of variance after pooling, the timing of pooling datasets, the best number of datasets to impute, the best method for handling imputations, at a minimum. Practical guidelines for methods that more efficiently and robustly account for the incompleteness of data within estimation methods based on the propensity score are needed, but research in this area is ongoing.

## Appendix 3 – Causal Inference

In the context of this work, the identifying assumptions required to draw causal inferences are: i) positivity, which implies that no individual has a probability of 1 of receiving Bolsa Familia conditional on their confounders, ii) consistency, which implies that different variations of receiving Bolsa Familia do not have different effects on TB outcomes, and iii) conditional exchangeability, which implies that there is no residual confounding.

It is broadly plausible that these assumptions hold. As demonstrated in Figure 3, Bolsa Familia is not deterministically assigned in practice, even to those families with no income. There are non-recipients of Bolsa Familia at a range of income levels below the income threshold used to determine eligibility so it is plausible that there is no individual with a probability of 1 receiving Bolsa Familia.

The exposure was specified as receiving any amount of Bolsa Familia for any amount of time. There are likely to be few variations in receipt of this variable when conceived in this manner, so consistency is plausible. However, future work should be conducted to determine how cure rate or other measures of TB burden vary with the amount of benefit received and the time for which it was received.

We can never rule out residual confounding, but the creation of a DAG to inform the analysis and the use of sensitivity analyses suggest the possibility is minimised, and thus conditional exchangeability is plausible. However, there remain some imbalances on state, and balance was not substantially improved for variables such as HIV and DOT that may have an impact on TB cure rate - the importance of these factors warrants further investigation.

A potential limitation to drawing causal conclusions is the non-interference assumption, which in this context assumes that the exposure received by one individual does not affect the outcome of the other. The results of this study suggest that the size of effect found may be too large to ignore this assumption and work should be undertaken to investigate the effect of social protection on TB transmission. Another potential violation of this assumption is that BFP increases the probability of cure not only in recipients but also in other cases through neighbourhood effects of the cash transfer.

We conclude that broadly these identifying assumptions are plausible, but refrain from drawing conclusions about causality given the interference limitations outlined above. The circumstances under which causal inferences can be drawn with interference is an area of ongoing research [30].

Authors’ Contributions
DJC, DB, and RhD conceived and designed the study. DJC analysed and interpreted data. DJC and DB drafted the manuscript. All authors provided intellectual contributions, reviewed, and approved the final version of the report.

